# Nuclear size-regulated emergence of topological packing order on growing human lung alveolospheres

**DOI:** 10.1101/2024.04.17.589951

**Authors:** Wenhui Tang, Jessie Huang, Adrian F. Pegoraro, James H. Zhang, Yiwen Tang, Dapeng Bi, Darrell N. Kotton, Ming Guo

**Affiliations:** Department of Mechanical Engineering, Massachusetts Institute of Technology, Cambridge, MA, 02139, USA; Center for Regenerative Medicine of Boston University and Boston Medical Center, Boston, MA 02118, USA; The Pulmonary Center and Department of Medicine, Boston University School of Medicine, Boston, MA 02118, USA; Metrology Research Centre, National Research Council Canada, Ottawa, Ontario, Canada; Department of Physics, Northeastern University, MA 02115, USA

## Abstract

Within multicellular living systems, cells coordinate their positions with spatiotemporal accuracy to form various structures, setting the clock to control developmental processes and trigger maturation. These arrangements can be regulated by tissue topology, biochemical cues, as well as mechanical perturbations. However, the fundamental rules of how local cell packing order is regulated in forming three-dimensional (3D) multicellular architectures remain unclear. Furthermore, how cellular coordination evolves during developmental processes, and whether this cell patterning behavior is indicative of more complex biological functions, is largely unknown. Here, using human lung alveolospheres as a model system, by combining experiments and numerical simulations, we find that, surprisingly, cell packing behavior on alveolospheres resembles hard-disk packing but with increased randomness; the stiffer cell nuclei act as the ‘hard disks’ surrounded by deformable cell bodies. Interestingly, we observe the emergence of topological packing order during alveolosphere growth, as a result of increasing nucleus-to-cell size ratio. Specifically, we find more hexagon-concentrated cellular packing with increasing bond orientational order, indicating a topological gas-to-liquid transition. Additionally, by osmotically changing the compactness of cells on alveolospheres, we observe that the variations in packing order align with the change of nucleus-to-cell size ratio. Together, our findings reveal the underlying rules of cell coordination and topological phases during human lung alveolosphere growth. These static packing characteristics are consistent with cell dynamics, together suggesting that better cellular packing stabilizes local cell neighborhoods and may regulate more complex biological functions such as organ development and cellular maturation.

## Main

The surfaces of three-dimensional (3D) multicellular systems, such as blood vessels, intestines, and pulmonary alveoli, are densely packed with cells. How cells coordinate to optimize their relative positions and nearest neighbor order is informative in various physiological and pathological processes^1–5^. For example, cell shapes change drastically during the extension of *Drosophila* germband epithelium^4^; in breast cancer clusters, cells on the boundary become larger and more elongated when forming invasive protrusions^5^. Moreover, cell nearest neighbor order changes in conjunction with the phase of tissues during jamming or glass transitions in a variety of simulation and experimental studies^2,3,6^. Despite these findings, it remains challenging to determine how tiling of cells affects structure formation and even biological functions such as maturation and invasion. Unlike monodispersed colloidal systems, packing in living tissues is highly disordered as cells are soft and variable in size. In addition, tissue and organ surfaces are naturally curved in 3D with non-zero Gaussian curvatures^7–11^, which adds another level of complexity to the packing problem. Although recent studies have shown that cell and tissue morphologies can be affected by a variety of factors, such as cell density^12–15^, curvature^13^, nuclear shape^13,15^ and mechanical forces^14^, it still remains unclear how soft cells tile a curved tissue surface and how cell packing is regulated, particularly during growth and maturation of 3D living systems. Here, using human induced pluripotent stem cell (iPSC)-derived lung alveolospheres as a model system, we observe an emergence of topological order, specifically a topological gas-to-liquid transition, during the growth of these alveolospheres, regulated by the increasing nucleus-to-cell size ratio and population size. As cell packing order increases, these spherical epithelia become less dynamic and more stabilized, with potentially stronger extracellular adhesion. Our finding reveals the critical role of cell nuclear size in regulating cell packing during tissue development, and suggests the importance of topological phase changes in establishing tissue stability.

To study the topological order of multicellular packing on curved biological surfaces, we use a human lung alveolosphere system cultured in 3D in Matrigel (Fig. 1a); this system contains a monolayer of human iPSC-derived alveolar epithelial type II cells (iAT2s) and exhibits a spherical geometry^16^. These iAT2s express a global transcriptome and ultrastructure that resemble primary adult alveolar epithelial type 2 cells (AT2s), thus serving as an *in vitro* model of AT2-related human lung development. As these alveolospheres grow in size, the total cell number within an alveolosphere increases, thus allowing us to investigate packing and topological order of the constituted iAT2s on these evolving spherical biological surfaces. To visualize cells in alveolospheres, we transduce them with a lentiviral vector to stably express Green Fluorescent Protein fused to a Nuclear Localization Signal (GFP-NLS) allowing nuclear localization of green fluorescence. With this system we monitor individual cell positions and behavior in each sphere by imaging entire alveolospheres using confocal microscopy^17^. We then track the positions of individual cell nuclei in 3D for alveolospheres of different sizes.

**Figure 1.**
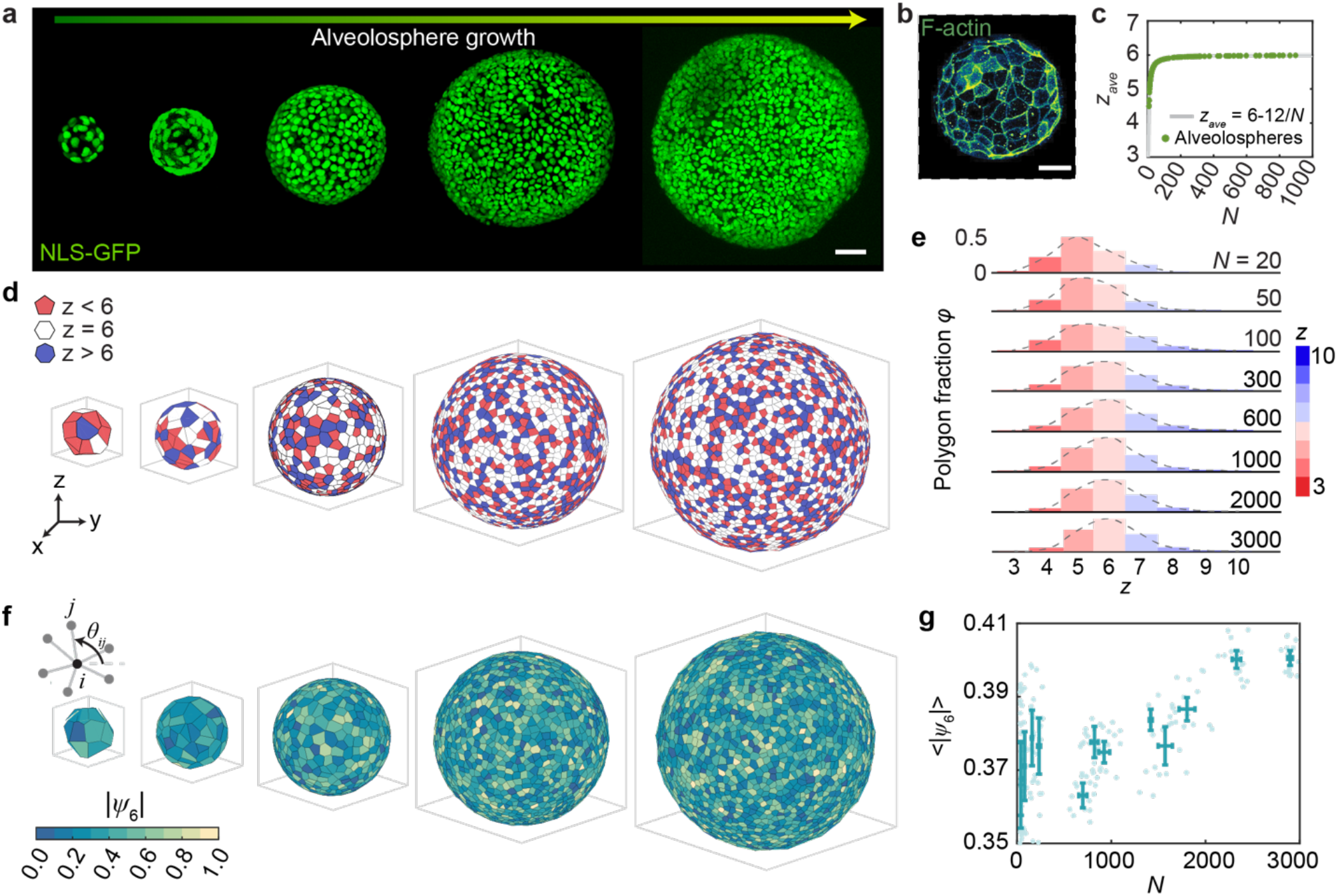
Emergence of topological order on growing human lung alveolospheres. **a**, Z-stack projections of GFP-NLS nuclei on representative growing 3D alveolospheres, scale bar: 50μm. **b**, F-actin staining of an alveolosphere showing the cell boundary, scale bar: 20μm. **c**, Average vertex number *z*_*ave*_ of all cells on each alveolosphere as a function of total cell number *N* (red dots) follows the theoretical curve *z*_*ave*_ = 6 −12/*N* (grey curve). **b**, Voronoi tessellation shows cell nearest neighbor order with three categories (z<6, z=6, z>6) for alveolospheres with increasing total cell number *N*. **e**, The distribution of different cell shapes showing that hexagonal cells become the predominant population as alveolospheres grow bigger. **f**, Bond orientational order |*ψ*_6_| for representative large alveolospheres. **g**, Average bond orientational order ⟨|*ψ*_6_|⟩ continues to increase as *N* increases, despite *z*_*ave*_ remaining almost constant. Mean ± std are shown.

## Results

### Emergence of topological order on growing human lung alveolospheres

Using the nuclear positions in alveolospheres (iPSC clone BU3 NGST, see Method), we obtain cell neighbor orders by performing Voronoi tessellation (Fig. 1d)^18,19^, which indicates the number of nearest neighbors for each cell. Compared to packing of monodispersed colloidal particles on spheres^7,9,20–22^, we find a significantly larger amount of nonhexagons with living biological cells on spheres (Fig. 1d). Nonetheless, cell packing still obeys simple topological principals controlling their structural organization—the average cell vertex number ⟨*z*_*i*_⟩ in alveolospheres aligns perfectly with the theoretical curve ⟨*z*_*i*_⟩ = 6 −12/*N* (*Supplementary Information*) ^23–25^, as shown in Fig. 1c. The cell packing on alveolospheres becomes more ordered as they grow bigger: hexagons gradually overcome pentagons as the predominant cell shape with increasing alveolosphere size *N* (Fig. 1e). Although the hexagon fraction plateaus at large *N* (*Supplementary Information*), we observe a slight increase in bond orientational order ⟨|*ψ*_6_|⟩^9,11^as *N* increases, suggesting a higher level of structural regularity and with cell shapes becoming closer to that of regular hexagons (Fig. 1f-g). Interestingly, if we compare the polygon fractions of packing of cells in alveolospheres to a random packing-on-sphere simulation (Method, Fig. S1), both plateau at large *N*, but alveolosphere packing has a higher ratio of hexagons than random packing (Fig. S1 inset). This difference in nearest neighbor order suggests that cell packing on alveolospheres is not random, and the arrangement of cells in spherical epithelia is influenced by factors beyond just topological constraint.

### Shortest nuclear dimension aligns with the minimum cell-cell distance and regulates nearest neighbor order on lung alveolospheres

To investigate what determines the distribution of nearest neighbor order for large systems, we perform a simulation to generate *N* particles on a unit sphere and limit the distance between neighbor particles to be no less than a minimum distance (*D*_*P*_)_*min*_ (Fig. 2a), which sets their effective diameter. To allow comparison between different total particle number *N*, as well as results from simulation and experiments, this minimum distance is nondimensionalized by the average occupied length *L*_ave_ for each particle and is defined as

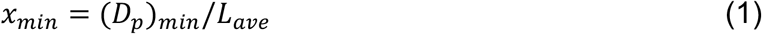

Where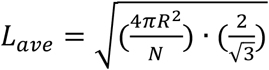 is calculated based on the particle average area with each *p*article treated as a hexagon. *x*_*min*_ also sets the area density of the effective hard-disk *p*acking, which is given by 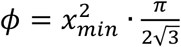(Supplementary Information). In simulation, when *x*_*min*_ varies from 0 to 0.7 for a certain system size *N*=800, we observe that the particle packing becomes more ordered as the minimum particle distance increases (Fig. 2b); hexagons (z=6) gradually become the predominant population and the fraction of polygons with both z>6 and z<6 decrease (Fig. 2c).

**Figure 2.**
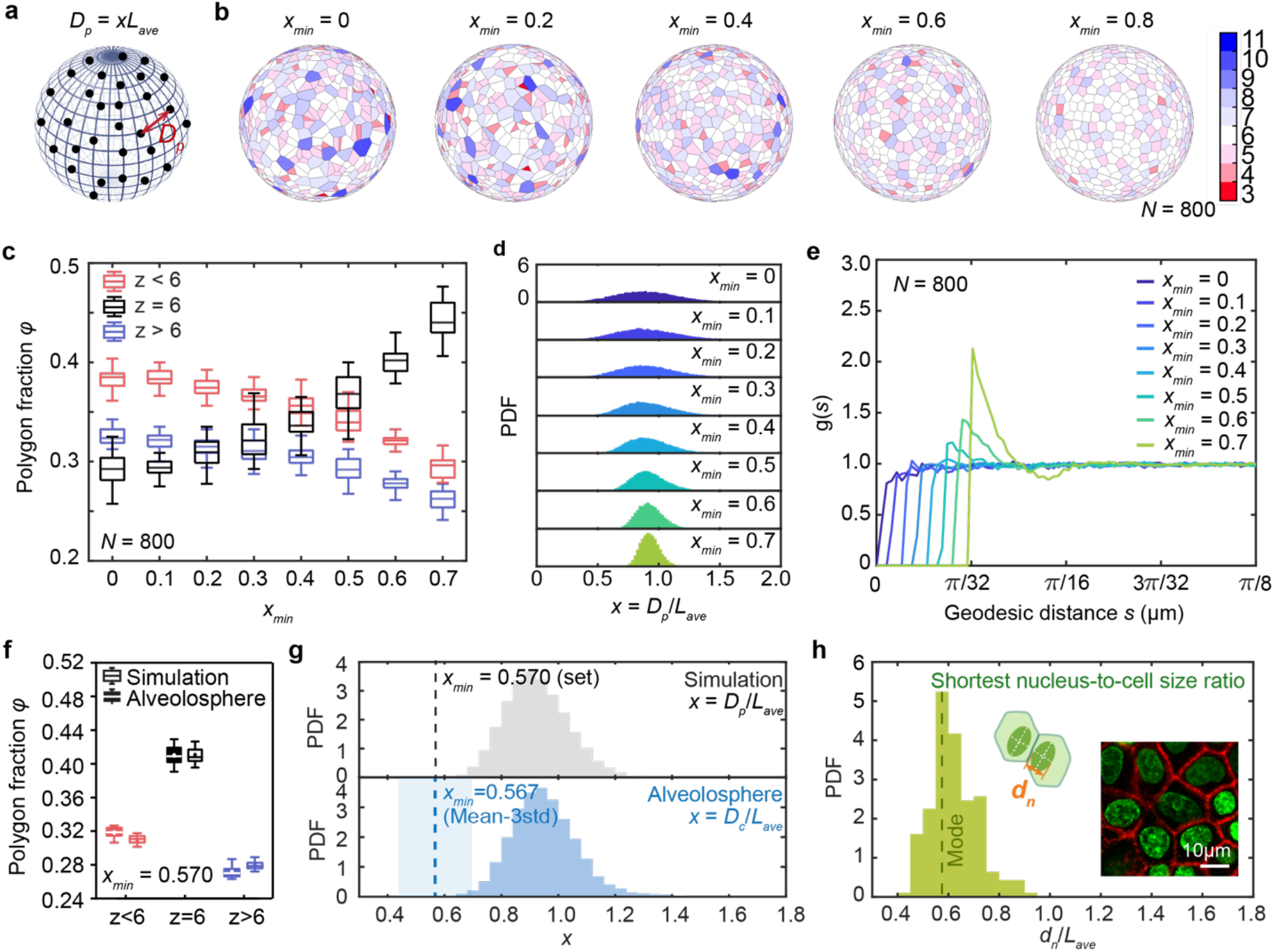
Nucleus to cell size ratio regulates nearest neighbor order on alveolospheres. **a**, Schematics of particles-on-sphere simulation, with the distance between particles defined as *D*_*P*_ = *x* · L_*ave*_, where *L*_*ave*_ is the average particle size and *x* is a ratio. **b**, Increasing *x*_*min*_ in simulation results in a higher fraction of hexagons and more ordered packing on a unit sphere. *N*=800. **c**, Quantification of polygon fractions shows that as *x*_*min*_ increases, the proportion of *z*=6 increases while the proportion of *z*<6 and *z*>6 decreases. Each box plot is from 30 tests of simulation. **d**, Varying *x*_*min*_ changes the distribution of nondimensionalized particle-particle distance *D*_*P*_/*L*_*ave*_ on spheres. **e**, Radial distribution function *g*(*s*) as a function of geodesic length *s* for different *x*_*min*_ when *N* = 800 on a unit sphere. Each curve is the average of 30 data sets. **f**, Polygon fractions for alveolospheres compared to particle simulations for *x*_*min*_ = 0.570 selected to best match. **g**, Probability density function (PDF) of *x* in alveolospheres agrees well with simulation when *x*_*min*_ = 0.570. Blue shaded region marks the region of (Mean−3std)±std, which is 0.567±0.129. **h**, Probability density function (PDF) of the shortest nucleus-cell size ratio for alveolosphere. The mode of the distribution is marked as the dashed line as 0.575. Inset: cell nuclei and cell boundary showing by F-actin. Scale bar: 10μ*m*.

To investigate how *x*_*min*_ regulates the short-range order on a sphere, we examine the particle-particle distance distribution under different *x*_*min*_ . The distribution of particle-particle distance becomes narrower as *x*_*min*_ increases (Fig. 2d), and this more uniform particle-particle distance yields a more ordered particle packing, which is consistent with recent study^26^. When comparing simulation results to experiments, we find that when *x*_*min*_ ≈ 0.57, the nearest neighbor order compositions (*z* < 6, *z* = 6, *z* > 6) are similar as to those observed in alveolospheres with the same system size (Fig. 2f, Fig. S7a). This

suggests that a limit of cell-cell distance also regulates the nearest neighbor order in alveolospheres. To further explore this, we plot the distribution of the nondimensional cell-cell distance *x* = *D*_0_/*L*_*ave*_, where *D*_*c*_ is cell-cell distance (Fig. 2g). Interestingly, we find that the histogram of *x* in alveolospheres resembles that from simulation with a preset *x*_*min*_ = 0.57 (Fig. 2g). These results are consistent with the existence of a minimum nondimensional cell-cell distance *x*_*min*_ = 0.57 where this *x*_*min*_ regulates the packing order on this alveolosphere. One estimate of this experimental *x*_*min*_ is to take the Mean-3std from the distribution *x* on the alveolosphere which we find is 0.567±0.129. To test if an experimental *x*_*min*_ is a universal feature of alveolospheres, we perform the same analysis on alveolospheres derived from another donor (iPSC clone SPC2-ST-B2^26^; see Methods) that naturally have smaller nuclear size and confirm that the cell packing is determined by minimum cell-cell distance as well (Fig. S3). These comparisons together show that the nearest neighbor order is regulated by the minimum cell-cell distance in human lung alveolospheres.

Within epithelia, cells cannot get infinitely close due to physical limitation. Given that the nucleus is the largest organelle in eukaryotic cells and is considerably stiffer than other cellular structures^25^, we wonder if cell nuclear size plays a major role in regulating the cell-cell distance and coordinating the nearest neighbor order^15^. In alveolosphere systems, cell nuclear shape is not circular but elliptical (Fig. S2), therefore the closest cell-cell distance is more likely to be limited by the shortest nuclear dimension *d*_*n*_ of the ellipse projected on the surface perpendicular to cell height (Fig. 2h, inset). When we measure

*d*_*n*_ for individual cells on alveolospheres, we can again determine a nondimensionalized representation by dividing by the average cell size *L*_*ave*_. This allows a comparison of this nondimensional shortest nuclear dimension *d*_*n*_/*L*_*ave*_ with the nondimensional cell-cell distance *x*, in particular its minimum value *x*_*min*_ (Fig. 2g-h). We find that *d*_*n*_/*L*_*ave*_ has a distribution aligning well with *x*_*min*_ (Fig. 2h), indicating that the nuclear size sets the minimum cell-cell distance. Furthermore, results of alveolospheres from a different donor (iPSC clone SPC2-ST-B2^26^; see Methods) that naturally have a smaller nuclear size confirm this finding as well (Fig. S3). To further investigate the idea that cell nuclear size sets the minimum cell-cell distance, and thus coordinates cell packing order on spherical surfaces, we apply hyper-osmotic and hypo-osmotic perturbations to change cell nuclear size by supplementing culture medium with 1.5% PEG300 and 20% DI water, respectively (Fig. 3a). We observe an increase in *d*_*n*_/*L*_*ave*_ under hyper-osmotic pressure and a decrease in *d*_*n*_/*L*_*ave*_ under hypo-osmotic pressure (Fig. 3c, e, Fig. S8); consistently, we find more ordered packing under osmotic compression (Fig. 3b-c), and less ordered packing under osmotic swelling (Fig. 3d-e). Together, these results confirm that the relative cell nuclear size regulates the topological packing on multicellular spheres, through limiting the minimum cell-cell distance.

**Figure 3.**
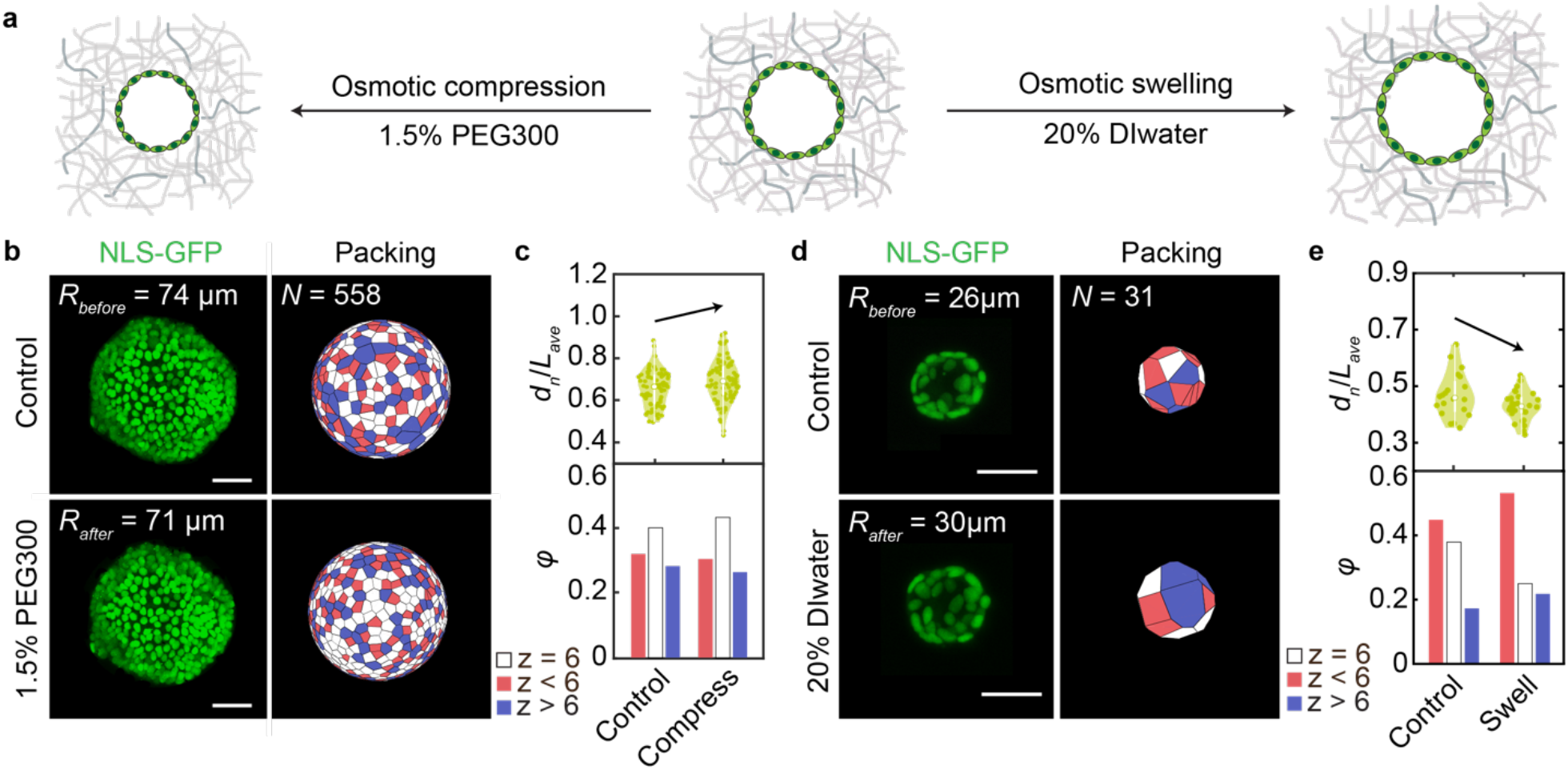
Perturbing the nucleus/cell size ratio changes cell packing in lung alveolospheres. **a**, Schematics of osmotic compression (add 1.5% PEG300) and swelling (add 20% DI water) to perturb alveolospheres. **b**, Osmotic compression squeezes the alveolospheres, causing increasing hexagon concentration in alveolospheres. Scale bars: 50 μ*m*. **c**, Polygon fraction and smallest nuclei ratio *d*_*n*_/*L*_*ave*_ evolution before and after osmotic compression. **d**, Osmotic swelling decreases hexagon concentration in alveolospheres. Scale bars: 50 μ*m*. **e**, Polygon fraction and smallest nuclei ratio *d*_*n*_/*L*_*ave*_ evolution before and after osmotic swelling.To understand how cell nuclear size dynamically regulates cellular organization on 3D curved surfaces during lung alveolosphere development, we measure the dimensionless shortest nuclear dimension *d*_*n*_/*L*_*ave*_ for individual cells on alveolospheres during their growth. Remarkably, we find that *d*_*n*_/*L*_*ave*_ increases during growth and seems to reach a plateau at large *N* (*N* > 500) (Fig. 4a). While both the average cell size and nuclear size decrease, the former decreases faster, resulting in an increasing ratio *d*_*n*_/*L*_*ave*_ as alveolospheres grow bigger and density increases (Fig. 4a inset). To understand the evolution of polygon fractions during alveolosphere growth, we compare cell shape fractions of the series of alveolospheres to simulations; in simulation, we set particle number *N* as total cell number and *x*_*min*_ as the average nucleus-to-cell size ratio *d*_*n*_/*L*_*ave*_which are experimentally measured in each alveolosphere. The experimental results are shown as semi-transparent dots and the corresponding simulation results are shown as solid symbols in Fig. 4b; we find that our simulations successfully replicate the measured cell shape fractions during alveolosphere growth. This again confirms that the total cell number and nucleus-to-cell size ratio regulate cell packing topology during human lung alveolospheres growth.

### Topological gas-to-liquid transition during alveolospheres growth

In the particle-on-sphere simulation, we examine the radial distribution function *g*(*s*) for systems with different minimum dimensionless cell-cell distance *x*_*min*_ (Fig. 2e). Interestingly, when the particles are randomly packed (*x*_*min*_ = 0), *g*(*s*) has similar features as an ideal gas without any apparent peaks, indicating there is no interaction between particles; when *x*_*min*_ becomes large (*x*_*min*_ = 0.7), *g*(*s*) develops the structure of liquid with a small number of peaks and valleys at the small distances, indicating short-range interaction between particles^27–32^. Thus, the radial distribution function captures a topological gas-to-liquid phase transition as the dimensionless minimum distance *x*_*min*_increases (Fig. 2e). To quantitatively study this topological phase transition behavior, we define a critical transition *x*_*c*_ as the *x*_*min*_ when a first peak in *g*(*s*) arises; this indicates that the interaction between nearest neighbors becomes significant (Method, *Supplementary Information*). Therefore, for different system size *N* from small (*N*=20) to large (*N*=1000), we can obtain a series of *x*_*c*_ forming a gas-liquid phase boundary line (Fig. S4). To correlate this emergent topological phase transition with the packing order, we plot a ternary phase diagram showing the coexistence of three nearest neighbor orders: *z* < 6, *z* = 6, and *z* > 6 (Fig. 4d and Fig. S5) and the corresponding *x*_*min*_ and system size *N* values. Each point in the ternary diagram corresponds to the polygon composition at a certain system size *N* and *x*_*min*_. Fig. S5 visualizes the contour lines of constant *N* with varying *x*_*min*_; from the bottom left to middle of the triangle, *N* increases. This agrees with the observation that when *N* is small, there are more cells with *z* < 6 and when *N* is large, there are more cells with *z* = 6 and *z* > 6 (Fig. 1d). Fig. 4d visualizes the contour lines of constant *x*_*min*_ with varying *N* and filled with colors in between (colors are labeled according to the values of *x*_*min*_ as shown in the caption); from left to bottom right *x*_*min*_ increases, showing that when *x*_*min*_increases there are more hexagons. The gas-liquid phase boundary that we obtained from *g*(*s*) with Fig. S4 can thus be plotted on the ternary phase diagram (dashed line in Fig.4d).

**Figure 4.**
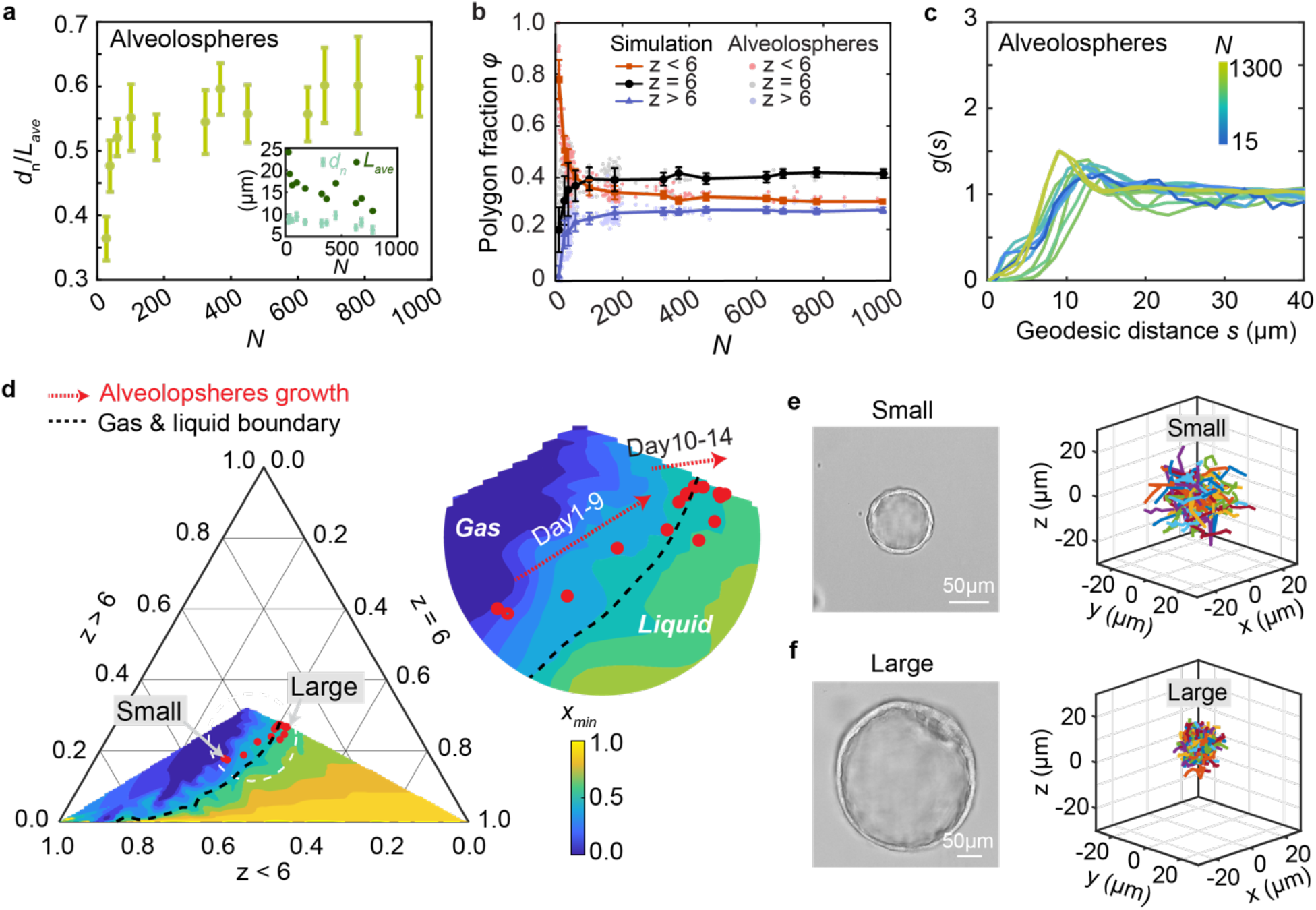
Increasing nuclear size ratio, a topological gas-to-liquid transition, and less dynamic cell migration as alveolospheres grow bigger. **a**, Evolution of smallest nucleus-to-cell size ratio *d*_*n*_/*L*_*ave*_ as a function of the total cell number *N* on alveolospheres. Inset: the separate trends of the smallest nuclei dimension *d*_*n*_ and average cell size *L*_*ave*_. **b**, Simulation using the experimentally measured *N* and *x*_*min*_ = d_*n*_/*L*_*ave*_ reproduces the nearest neighbor order fractions in a series of growing alveolospheres. Semi-transparent dots: alveolospheres data; solid lines: simulation. **c**, Radial distribution function *g*(*s*) of geodesic distance *s* for alveolospheres with different sizes. **d**, Ternary phase diagram of *z* < 6, *z* = 6, *z* > 6 visualized as a function of *x*_*min*_. Background color represents *x*_*min*_ from simulation. Red points show the alveolosphere data. Dashed line marks the gas-liquid phase boundary defined from the coordination number at different *N* (Supplementary *Information*). **e**, Cell trajectories for 50 randomly selected cells within a small alveolosphere (*R* = 30 μ*m*) for 10 continuous frames with time interval 15 mins each. **f**, Cell trajectories for 50 randomly selected cells within the largest alveolosphere (*R* = 160μ*m*) for 10 continuous frames. Time interval of each frame is 15 mins.

As *x*_*min*_ in the simulation has been confirmed to align well with *d*_*n*_/*L*_*ave*_ experimentally measured in alveolospheres, we thus overlay the nearest neighbor order compositions (z< 6, *z* = 6, and *z* > 6) of the growing alveolospheres on the ternary phase diagram obtained from simulation (Fig. 4d). Each red dot in Fig. 4d represents a nearest neighbor composition of a certain alveolosphere with a system size *N* and nucleus-to-cell size ratio *d*_*n*_/*L*_*ave*_ . As alveolospheres grow bigger, we have shown that there is an increasing number of hexagons; interestingly, we also notice that the polygon compositions of growing alveolospheres pass across the defined gas-liquid phase boundary (Fig. 4d). The idea that a phase transition is occurring as the alveolosphere develops is consistent with changes in the radial distribution function *g*(*s*): as alveolospheres become bigger, they exhibit significant short-range interactions among neighboring cells (Fig. 4c). This finding suggests that the growing alveolospheres undergoes a topological gas-to-liquid phase transition. The increase of system size *N* causes an obvious increase of hexagon fractions in the range of smaller-sized systems (approximately *N*<200); in larger systems (approximately *N*>200), the increased topological order is mainly driven by the naturally increasing *d*_*n*_/*L*_*ave*_ as this alveolosphere system grows bigger and cells becomes denser. To further test this, we manipulate the nucleus to cell size ratio by applying hyper-osmotic pressure or hypo-osmotic pressure, respectively; we indeed find that the multicellular state can be pushed from the gas-liquid boundary either to a more liquid-like state or to a more gas-like state (Fig. S6). Moreover, a material phase transition should also impact dynamic behavior of the constituent particles. To test this behavior, we examine the cell dynamics of the large and the small alveolospheres on two sides of the phase boundary and find that cells are less motile in the fluid phase compared to gas phase (Fig. 4e-f); this is also in agreement with our previous observations of less dynamic multicellular flow on larger alveolospheres^17^.

## discussion

In conclusion, using human iPSC-derived lung alveolospheres and particles-on-sphere simulation, we investigate how cells tile 3D curved tissue surfaces, and discover that cell packing topology is highly regulated by both nucleus/cell size ratio and total cell number within the system. In addition, we find that nucleus/cell size ratio coordinates cell packing and drives the topological phase evolution of the curved cell epithelia. With an increasing nucleus/cell size ratio in growing alveolospheres, we observe a progressive emergence of short-range order, increasing number of cells with six nearest neighbors, and a coupled topological gas-to-liquid transition. This increase in nucleus/cell size ratio has also been observed during the development of pre-implanted mouse embryos from single cell to blastocyst^33^ and the starfish embryos^34^, suggesting that this might be a general phenomenon during morphogenesis. In fact, the nucleus is considered the control center in eukaryotic cells which contains the genetic information of life. Being able to regulate its relative size to the cell size is important to control the intranuclear environment (e.g., the degree of crowding) and thus a myriad of biological functions^33–37^. Moreover, since the stiff cell nucleus accounts for a larger portion of the cell during morphogenesis, it can potentially contribute to an increase in epithelial rigidity that facilitates stabilization of tissue geometry and thus maturation. Indeed, as these iPSC-derived alveolospheres grow bigger, cell migration slows down, resulting in a more stabilized cell neighborhood; this potentially can promote a more stable cell-cell communication, influencing higher-level biological functions. Finally, the basic rule of how soft living cells tile curved tissue surfaces may have a broad implication to designing soft robotics, wearable sensors, as well as metamaterials.

## Methods

### iAT2 3D culture

Human induced pluripotent stem cell-derived alveolar epithelial type II cells (iAT2s) were generated via directed differentiation as previously described^26,38,39^. Two iPSC lines were employed as indicated in the text: clone SPC2-ST-B2 and clone BU3 NGST. After differentiation into iAT2s, cells were maintained in 3D Matrigel (Corning No. 354230) as “alveolospheres” at 400 cells/μl Matrigel, and passaged every 10-14 days, as published^26,38,39^. A lentiviral vector with constitutively and ubiquitously active long EF1a promoter (EF1aL) driving expression of nuclear localized GFP (nlsGFP) was engineered by cloning a nuclear localizing sequence (NLS) in front of GFP. The resulting construct (pHAGE-EF1aL-nlsGFP-W) with full plasmid map and sequence are available from Addgene, plasmid #126688. Lentiviral infection of iAT2s was performed in suspension culture at a multiplicity of infection of 20, as previously detailed^39^, and NLS-GFP expressing cells were purified at a subsequent culture passage through fluorescence activated cell sorting and replated for serial sphere passaging in 3D culture. For imaging, NLS-GFP-expressing iAT2s were seeded in 3D Matrigel droplets in 35mm glass-bottom petri dishes at 100-200 cells/μL. These cells grow, proliferate, and self-organize into alveolospheres in 3D conditions.

### Image acquisition

To visualize the cell nuclei, cells with green fluorescent protein fused to a nuclear localization signal (NLS-GFP) were used. During imaging, alveolospheres were cultured in 35 mm petri dish in a customized incubator (5% CO_2_, 37°C, 95% humidity) on a confocal microscope (Leica, TCL SP8). A 25 × /0.95 numerical aperture water objective and a 63×/1.20 numerical aperture water objective were both used for imaging. Confocal microscopy recorded cells nuclear positions in 3D. Cell nuclear positions were then tracked using *Trackmate* plugin in *ImageJ*. Immunofluorescence images were taken by 63×/1.20NA water objective.

### Random packing-on-sphere simulation

In the simulation of randomly putting N particles on a unit sphere, the spherical coordinates (*θ, φ, R*) are randomly generated using uniformly distributed *rand* function in MATLAB. Specifically, *θ* = *π* · *rand* and *φ* = 2*π* · *rand*, and the spherical coordinate is converted to Cartesian coordinates for Voronoi tessellation.

### Bond orientational order |*ψ*_6_|

Bond orientational order at each cell with position *r*_*i*_ is calculated following the equation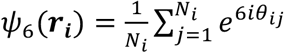, where *N*_*i*_ is the coordination number of the *i*^th^ cell, and *θ*_*ij*_ is the angle between the line connecting cell *i* to its *j*^th^ neighbor and an arbitrary reference axis. On a flat surface, the largest ⟨|*ψ*_6_|⟩ can reach 1; however, on a sphere, ⟨|*ψ*_6_|⟩ reaches a plateau of ℼ0.87.^9^ Bond orientational order MATLAB code was adapted from here^9^: https://www.nature.com/articles/nature25468#additional-information.

### Determining the gas-liquid phase boundary

During simulations, we gradually increase *x*_*min*_ and define a gas-liquid phase boundary as the *x*_*min*_when a significant first peak starts to appear in radial distribution function. We define this critical transition point as *x*_*c*_. Specifically, we define the *x*_*c*_ as the *x*_*min*_at which point the integration of *g*(*s*) from 0 to the first valley equals 1,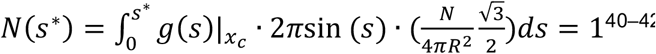, where *s*^*^ is the geodesic distance for the first valley.

### Osmotic perturbation

Osmotic compression is applied by adding 1.5v/v% polyethylene glycol 300 (PEG300) to isotonic culture medium. Osmotic swelling is applied by adding 2.0v/v% DI water to isotonic culture medium.

### The smallest nuclei dimension

As the shape of cell nuclei is not circular in alveolospheres, the smallest nuclei dimension is captured as the smallest dimension of all the directions for cell nuclei.

## Acknowledgment

We acknowledge the helpful discussion with David R. Nelson, Yiwei Li, and Roger D. Kamm. We also acknowledge the MIT *SuperCloud* and Lincoln Laboratory Supercomputing Center for providing HPC resources. W. T. thank the Mathworks Engineering Fellowship; M.G. thank National Institute of General Medical Sciences grant number 1R01GM140108 and Sloan Research Fellowship; D.N.K. acknowledge support from NIH grant numbers U01HL148692, U01HL134745, U01HL134766, R01HL095993, and N01 75N92020C00005.

## Contributions

W.T. and M.G. designed the experiments. M.G. supervised the project. W.T. and J.H. performed the experiments. W.T. did image processing and data analysis. J.H.Z., Y.T., D.B. and A.F.P. provided essential perspectives for data analysis. W.T., A.F.P., Y.T., J.H., D.N.K., D.B. and M.G. wrote the manuscript. All authors edited and approved the manuscript.

## Competing interests

The authors declare no conflict of interests.

## Data and materials availability

Data supporting the findings of this study are available within the article and its Supplementary Information files and from the corresponding authors upon request. MATLAB scripts used in this work are available from the corresponding authors upon reasonable request.

## Notes

### Competing Interest Statement

The authors have declared no competing interest.

