## Supplementary Information for "Nuclear size-regulated emergence of topological packing order on growing human lung alveolospheres"

**This file includes:**
Fig S1-S8
SI text
SI references

**Check the topological constraint of cell packing on spheres**

The nearest neighbor order for each cell is defined by its vertex number $z_i$, and the average cell vertex number on each alveolosphere is $\langle z_i \rangle$. In the theory in colloidal crystallization [1, 2], cells that do not have six vertexes are defined as topological defects, and the corresponding topological charge for each defect cell is defined as $q_i = 6 - z_i$. For a given system, we calculate the total topological charge $Q = \sum_{i=1}^{N}(6 - z_i)$, where $N$ is the total cell number on an alveolosphere [1, 2]. For a sphere, a total +12 charges are required by a classical theorem of Euler equating the total disclination charge to $6\chi$, where the Euler characteristic $\chi$ is 2 for the sphere [1-3].

**Plateau of polygon fractions at large *N***

All the polygon fractions reach plateaus when N is large, indicating that the population size N only strongly affects polygon fractions on small alveolospheres, while its effect is diluted when N is large. The dominance of N is evident with more variations in polygon fractions at small system size (FIG. S1). As we compare these results with a random particle-on-sphere simulation on a sphere, we find that although they both have plateaus at large *N*, the three compositions of cell nearest neighbor order (z<6, z=6, z>6) are significantly different (FIG. S1 inset). This difference in nearest neighbor order suggests that cell packing on spherical surfaces is not entirely random, and the topological constraint is not the only factor that decides cell arrangement in spherical epithelia.

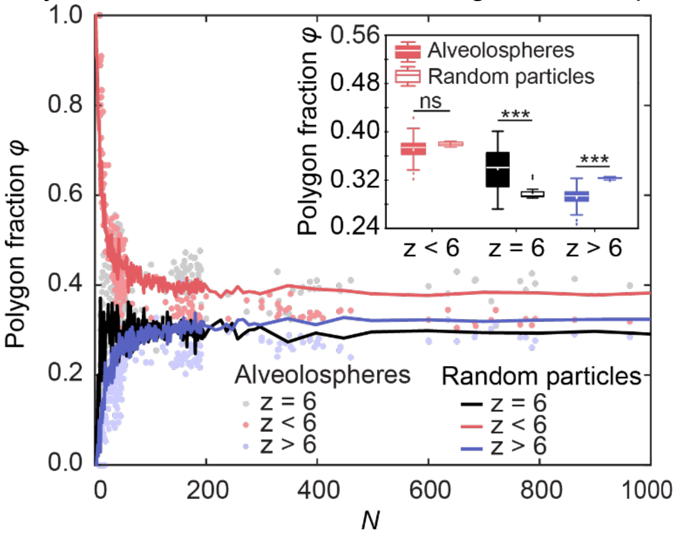


**Figure S1.** Comparison of the evolution of polygon fractions in growing lung alveolospheres and a random packing on sphere. Inset: box plot showing the three categories of polygon fractions (z>6, z=6, z<6) at the plateau region.

**Dimensional analysis of alveolosphere system**

In the alveolospheres system, at every observation at certain time, we have the following variables:

$$N, \rho, d_n$$

where $N$ is the cell number, $\rho = \frac{N}{4\pi R^2}$ is the particles packing density (where $R$ is the sphere radius), and $d_n$ is the nuclei length scale. Their SI units are:

| Variables | SI units |
|---|---|

| | |
|---|---|
| $N$ | - |
| $\rho$ | $L^{-2}$ |
| $d_n$ | $L$ |

Therefore, we form two $\Pi$ groups that can describe the system behavior:

$$N, d_n\sqrt{\rho}$$

**Compare $x_{min}$ with the hard sphere packing fraction $\phi$**

Assume $x_{min}$ corresponds to a hard sphere diameter $D_{p2\min} = x_{min}L_{ave}$ where the hard spheres cannot overlap, i.e. the distances between hard spheres must be larger than $D_{p2pmin}$. According to the definition,

$$\phi = \frac{\text{hard sphere area}}{\text{total surface area}}$$

we have

$$\phi = \frac{N\pi(\frac{D_{p2pmin}}{2})^2}{4\pi R^2} = \frac{N(\frac{D_{p2pmin}}{2})^2}{4R^2}$$

$$= \frac{N\left[x_{min} \cdot \sqrt{\left(\frac{4\pi R^2}{N}\right) \cdot \left(\frac{2}{\sqrt{3}}\right)}\right]^2}{16R^2}$$

$$= x_{min}^2 \cdot \frac{\pi}{2\sqrt{3}}$$

where $\phi$ is only a function of $x_{min}$. Therefore, when $x_{min}$ in range [0, 0.7], packing fraction is in a range of [0, 0.444]. This range is smaller compared to the maximum packing fraction for random packing of hard disks with unit size which is ~0.85.

**Particle-on-sphere simulation on unit spheres**

The particle-on-sphere simulation is performed using Random Sequential Addition (RSA) method on unit spheres. Basically, particles are randomly generated with spherical coordinates on a unit sphere one by one, and then the spherical coordinates are converted into Cartesian coordinates. The newly generated particle will be kept on the sphere if the distances between it and its nearest neighbors are within the set value:

$$(D_p)_{min} = x_{min} \cdot L_{ave}$$

Otherwise, it will be removed, and a new particle will be generated again at a random position on sphere. This process will loop until the total number of particles reaches to the set total system size *N*. When the loop ends, there will be *N* particles sitting on the sphere and the distance between any nearest neighbor particles is larger than $(D_p)_{min}$ that we set. To make comparison between different system sizes, we nondimensionalize the $(D_p)_{min}$ by average cell size $L_{ave}$

$$x_{min} = (D_p)_{min}/L_{ave}$$

And we vary $x_{min}$ for different cases in simulation.

**Radial distribution function $g(s)$**

The radial distribution function is calculated along the geodesic distance. The density variation is first calculated for each particle before taking an ensemble average,

$$g(s) = \frac{1}{N}\sum_{i=1}^{N} g_i(s)$$

Where N is the total cell number and $g_i(s)$ is the radial distribution function calculated for each cell using the following formula,

$$g_i(s) = \frac{\rho(s)}{\rho_{bulk}} \approx \frac{\frac{dn(s)}{2\pi r sin\theta \cdot d(s)}}{\frac{N}{4\pi r^2}}$$

where $dn(s)$ is the cell number found within a short geodesic distance range $d(s)$ at $s$.

**Determine the gas-liquid phase boundary using radial distribution function**

The gas-to-liquid phase boundary is determined using particles-on-sphere simulation. For a series of different system sizes *N*, we plot the radial distribution functions $g(s)$ with different $x_{min}$, as shown in Fig.S4. When $x_{min}$ is small, radial distribution function shows no peak, suggesting there is no interaction between particles like an idea gas. When $x_{min}$ is large, the radial distribution function shows short-range peak and valley, which is similar to is observed both experimentally and in simulations for liquids [4-9]. Given these two end points, there must be a transition from ideal gas to liquid in between them. We thus define a critical transition $x_c$ here as the $x_{min}$ when a significant first peak starts to appear in the radial distribution function as we gradually increase $x_{min}$. Although a single peak in radial distribution function is observed in hard sphere 'gas' simulations [10], here we use the appearance of first peak as a signature that the system is no longer ideal gas but starts to show significant particle-particle interaction. To have a quantitative identification, we define the $x_c$ as the point at which the integration of $g(s)$ from 0 to the first valley equals 1,

$$N(s^*) = \int_0^{s^*} g(s)|_{x_c} \cdot 2\pi \sin{(s)} \cdot (\frac{N}{4\pi R^2}\frac{\sqrt{3}}{2})ds = 1$$

suggesting there is one particle in the nearest neighbor, where $s^*$ is the geodesic distance when the first valley appears [11-13]. Then for each system size *N* we can find a critical $x_c$. The series of *N* and $x_c$ correspond to different nearest neighbor compositions on the ternary phase diagram, and we mark the self-defined phase boundary as a black dashed line (Fig.4d right).

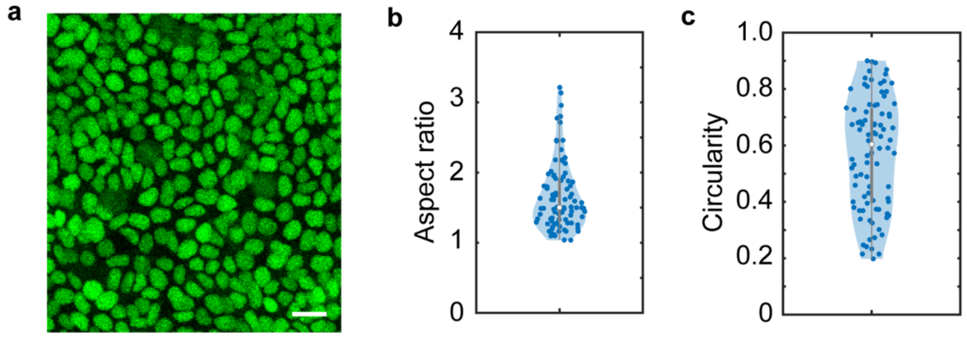

**Figure S2**. Cell nuclei in alveolospheres are not spherical but elliptical. **a**, Representative cell nuclei GFP-NLS showing the nucleus shape. Scale bar: 20 $\mu m$. **b**, Violin plot of aspect ratio for cell nuclei. **c**, Violin plot of circularity for cell nuclei.

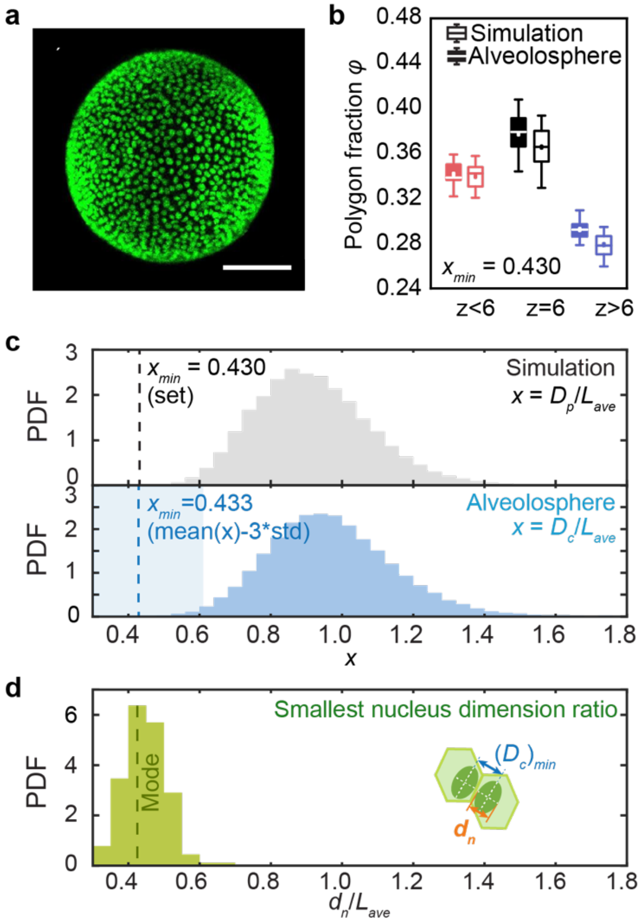


**Figure S3.** Alveolosphere (clone SPC2-B2) that naturally has a smaller nuclear-cell size ratio shows nuclear-cell size ratio aligns with the smallest cell-cell distance ratio and regulates packing fraction. a, NLS-GFP of a representative alveolosphere which has *N*~1200. Scale bar: 100 $\mu m$. **b**, Polygon fractions for alveolospheres compared to particle simulations for $x_{min} = 0.43$ selected to best match. **c**, Probability density functions (PDFs) of $x$ in alveolosphere agrees well with particle-particle distance when $x_{min} = 0.43$. Blue shade marks the region of (Mean-3std)$\pm$std$= 0.433 \pm 0.183$. **d**, Probability density functions (PDFs) of the shortest nuclei ratio $d_n/L_{ave}$ for alveolosphere. The mode of the distribution is label as dashed line in the middle of the bar at $d_n/L_{ave} = 0.425$.

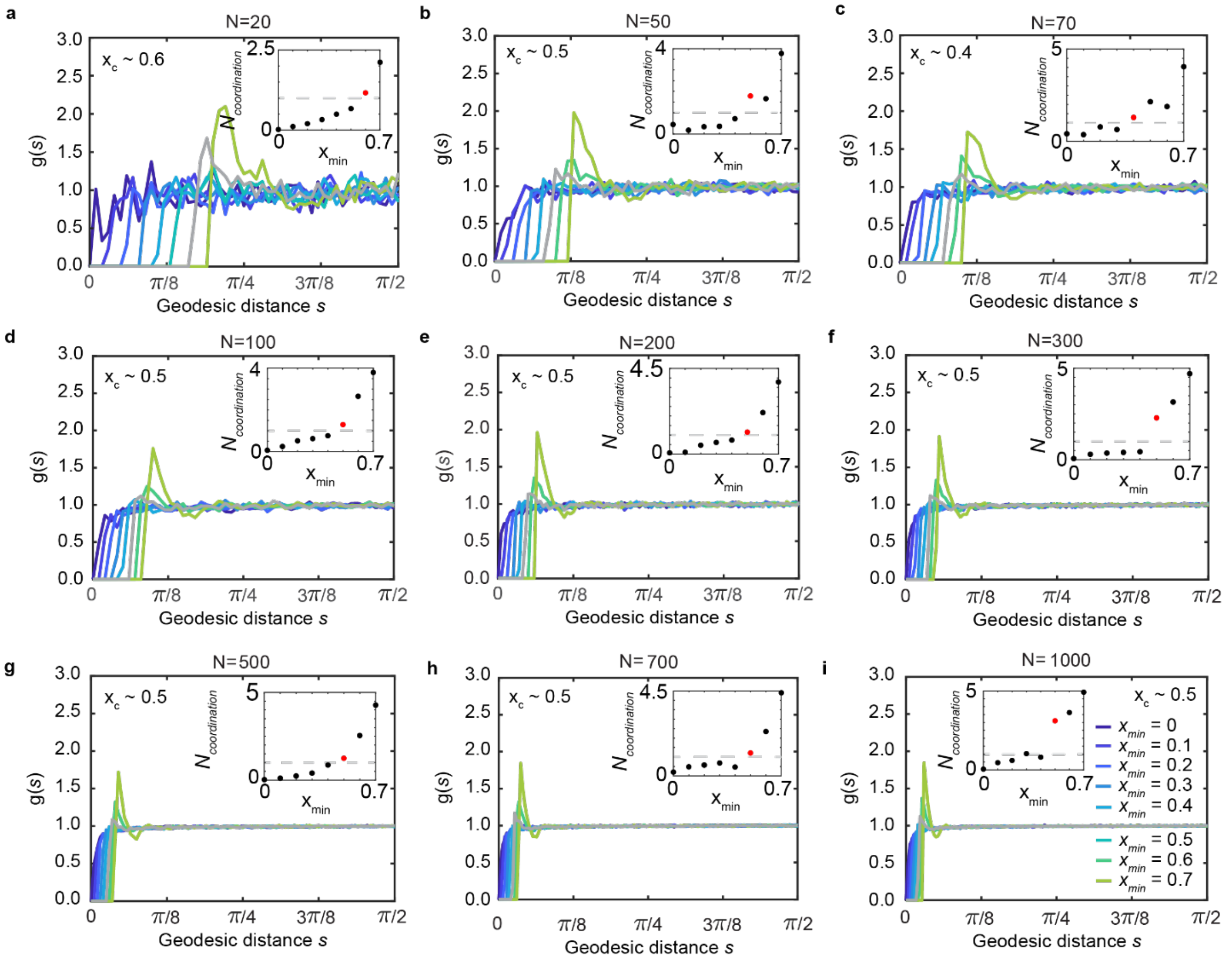


**Figure S4** Radial distribution function (RDF) and coordination number for different minimum particle-particle distance ratios $x_{min}$. Total particle number: **a-i**, *N*= 20, 50, 70, 100, 200, 300, 500, 700, 1000. Inset: coordination number as a function of $x_{min}$. From the radial distribution function, we can see a transition from gas to liquid as $x_{min}$ increases. We thus estimate $x_c$ for the series of *N* when the average coordination number starts to become larger than 1, as shown in grey RDF and red dots in the inset.

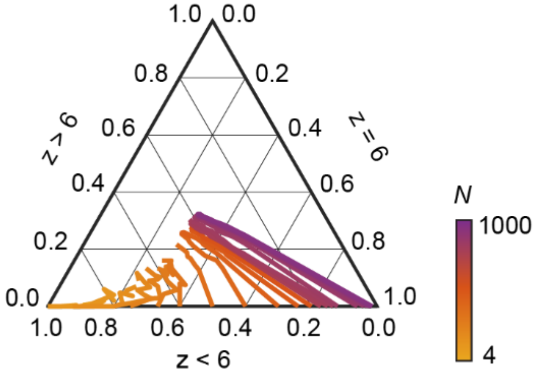


**Figure S5**. Ternary phase diagram of $z < 6, z = 6, z > 6$ as a function of $N$.

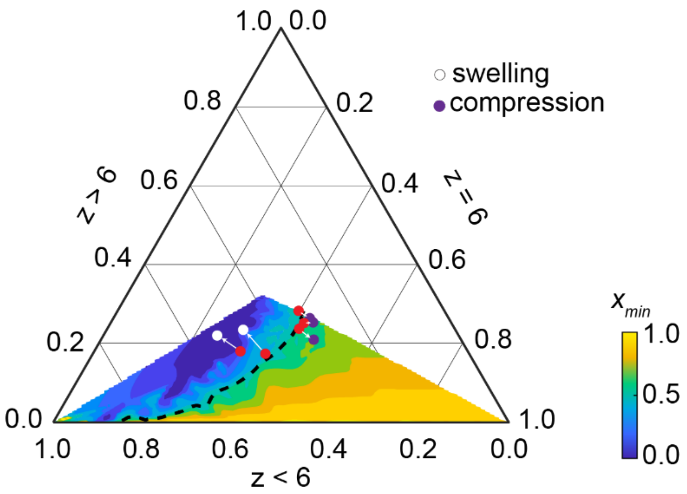


**Figure S6.** Ternary phase diagram showing the evolution of cell nearest neighbor order composition after osmotic compression (purple dots) and osmotic swelling (white dots).

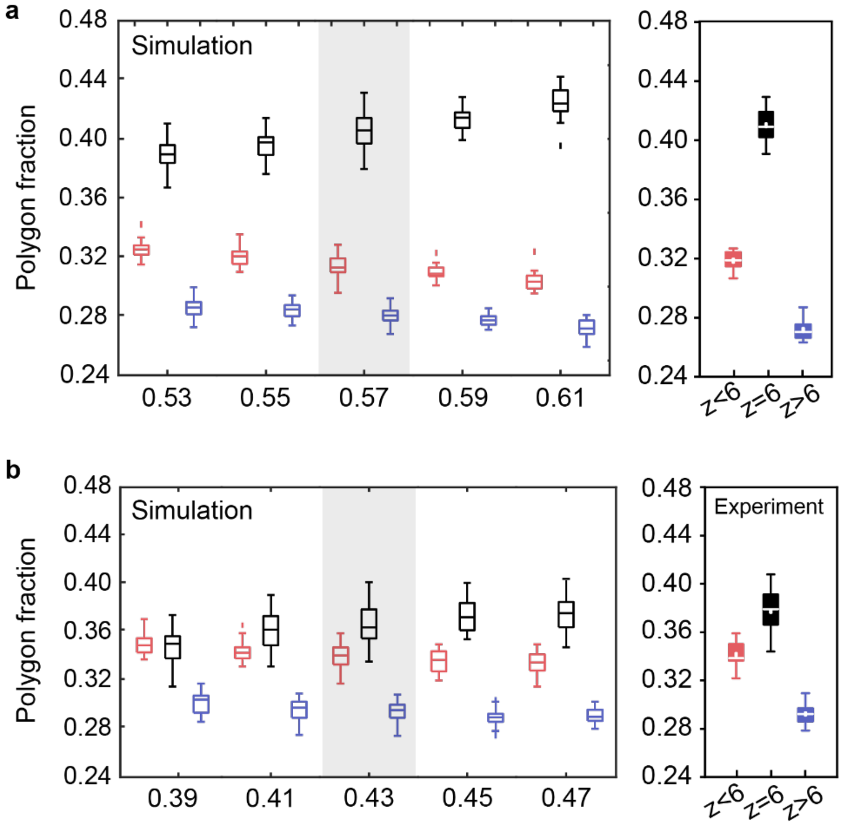


**Figure S7**. Find the $x_{min}$ in simulation that matches the most with experiment. a, Test for alveolosphere (clone BU3 NGST) with $x_{min} = 0.57$. Right: Simulation showing the polygon fractions with $x_{min} = 0.53, 0.55, 0.57, 0.59, 0.61$. Left: Polygon fractions in experiments. b, Test for alveolosphere (clone SPC2-ST-B2) with $x_{min} = 0.43$. Right: Simulation showing the polygon fractions with $x_{min} = 0.39, 0.41, 0.43, 0.45, 0.47$. Left: Polygon fractions in experiments

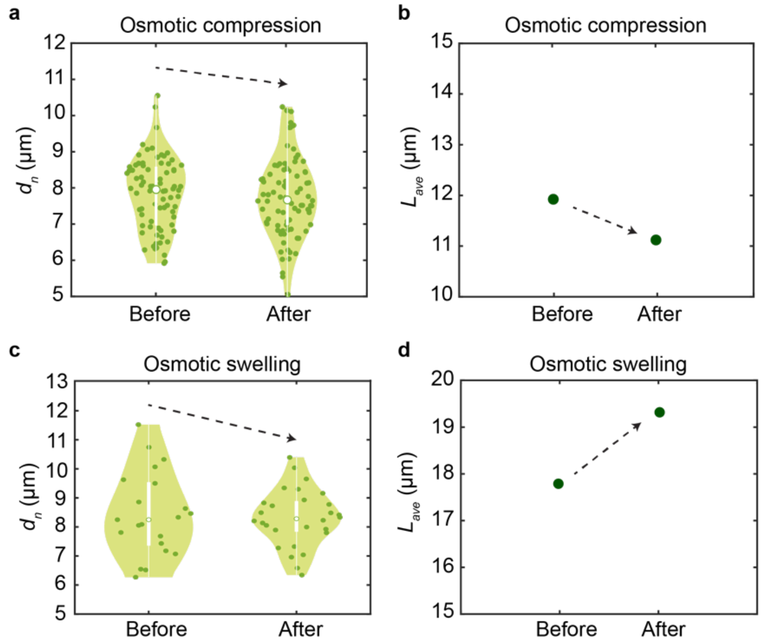


**Figure S8.** The change of $d_n$ and $L_{ave}$ during osmotic compression and swelling. After osmotic compression, $L_{ave}$ (calculated based on *R* and *N* before and after) decreases from 11.93$\mu m$ to 11.45$\mu m$. After osmotic swelling, $L_{ave}$ (calculated based on *R* and *N* before and after) increases from 17.79$\mu m$ to 20.53$\mu m$.
